# The metabolic mechanisms underlying zooplankton-derived dissolved organic matter’s chemical properties

**DOI:** 10.1101/2024.10.27.620474

**Authors:** Muhammad Firman Nuruddin, Ding He, Longjun Wu

## Abstract

Dissolved organic matter (DOM), the largest reservoir of organic material in the ocean, plays a crucial role in the global nutrient cycle and the microbial loop. While existing studies have documented significant DOM release by zooplankton, the chemo diversity and properties of this DOM, along with the physiological mechanisms influencing these characteristics in the environment, remain inadequately explored. We conducted zooplankton sampling followed by onboard DOM release experiments in heterogeneous estuarine-coastal water systems, followed by molecular characterization of the DOM using Fourier-transform ion cyclotron resonance mass spectrometry. Additionally, we analyzed zooplankton metabolic activities through meta-transcriptomics to elucidate the relationship between the chemical properties of the released DOM and the underlying physiological processes of zooplankton. Our findings reveal substantial variations in the molecular diversity of DOM released by zooplankton, more specifically lipids-like, protein-like, and unsaturated hydrocarbon-like between mesotrophic and eutrophic coastal zooplankton communities. We found strong correlations between chemical composition of the DOM and zooplankton gene functions associated with metabolism processes such as carbohydrate metabolism, nucleotide processing, energy production, and coenzyme metabolism. Furthermore, the modified aromaticity indexes of the released DOM are also highly associated with metabolism-related gene functions such as amino acid, carbohydrate metabolism, lipid, energy production, as well as glycan biosynthesis, indicating that zooplankton metabolic processes significantly influence DOM aromaticity. This study enhances our understanding of how organism’s metabolic processes shape the molecular characteristics of DOM they release, highlighting its implications for nutrient cycling in the environment.

## Introduction

Dissolved organic matter (DOM) constitutes the largest pool of organic material in the ocean, driving global biogeochemical cycles and sustaining microbial communities through the microbial loop^1,2^. Comprising a complex mixture of bioavailable and refractory compounds, DOM serves as energy source for microbial heterotrophs, modulates nutrient and trace metal bioavailability, and alters the optical characteristics of water bodies^3–6^. While autochthonous DOM derived from phytoplankton exudates has been extensively studied, it accounts for only ~50% of heterotrophic microbial carbon demand, suggesting critical gaps in identifying other key DOM sources within aquatic ecosystems^7–10^.

Metazoan zooplankton serves as a link between primary producers and microbial loops, mediating DOM transfer via excretion, sloppy feeding, and digestion^1,11–14^. Unlike phytoplankton-derived DOM—which predominantly contributes to relatively more refractory pools^15–17^—zooplankton releases labile components such as amino acids and protein, which can serve as important nutritional sources for both autotrophic and heterotrophic microbes^8,18,19^. By modulating the availability of these labile substrates, zooplankton indirectly regulate microbial community dynamics, drive shifts in nutrient remineralization rates and alter the efficiency of nutrients transfer through aquatic food webs. On one hand, despite substantial advancements in our understanding of metazoan zooplankton’s ecological functions, limited studies focus on characterized their derived DOM molecules composition^19,20^. On the other hand, although prior studies have focused on elucidating zooplankton gene expression under environmental heterogeneity^21–24^ but have not explained the relation of these processes to molecular-level of the released DOM dynamics, leaving a critical disconnect between organismal biology and biogeochemical outcomes. Therefore, elucidating the physiological processes (gene expression) of zooplankton together with the variations of their derived DOM is critical for capturing a holistic understanding of the animal’s role in biochemical cycle within aquatic ecosystems.

Advances in high-resolution mass spectrometry and community-level omics now enable unprecedented exploration of organism metabolic-DOM linkage. Techniques such as Fourier Transform Ion Cyclotron Resonance Mass Spectrometry (FTICR-MS) allows for the detailed characterization of DOM molecular formulae^30,31^, while meta transcriptomics enable the examination of organism including zooplankton gene expression which elaborates their physiological and metabolism processes at the community level^32^. Here, we integrate these tools to test the hypothesis that zooplankton metabolism gene expression variabilities govern the molecular diversity and functional traits of released DOM. Through spatially resolved sampling across trophic level gradients, eutrophic to mesotrophic in pearl river estuary and adjacent coastal regions, the second largest river in China, followed with DOM release experiments, we uncover relationships between zooplankton metabolic pathways—including amino acid, carbohydrate metabolism, lipid, energy production, as well as glycan biosynthesis and metabolism—and DOM chemical signatures. Our findings establish organism physiology as a key determinant of autochthonous marine DOM dynamics, advancing mechanistic understanding of animal physiological process role in marine biogeochemistry.

## Result and Discussion

### Zooplankton derived DOM overall variability

To characterize dissolved organic matter (DOM) produced by zooplankton physiological activity in heterogeneous coastal waters, we conducted controlled zooplankton release incubations targeting DOM from metabolic excretion and faecal pellet leaching. These experiments generated 15 DOM samples across seven sites spanning a trophic gradient from the eutrophic Pearl River Estuarine delta to adjacent mesotrophic coastal waters (Fig. 1a).

**Fig 1.**
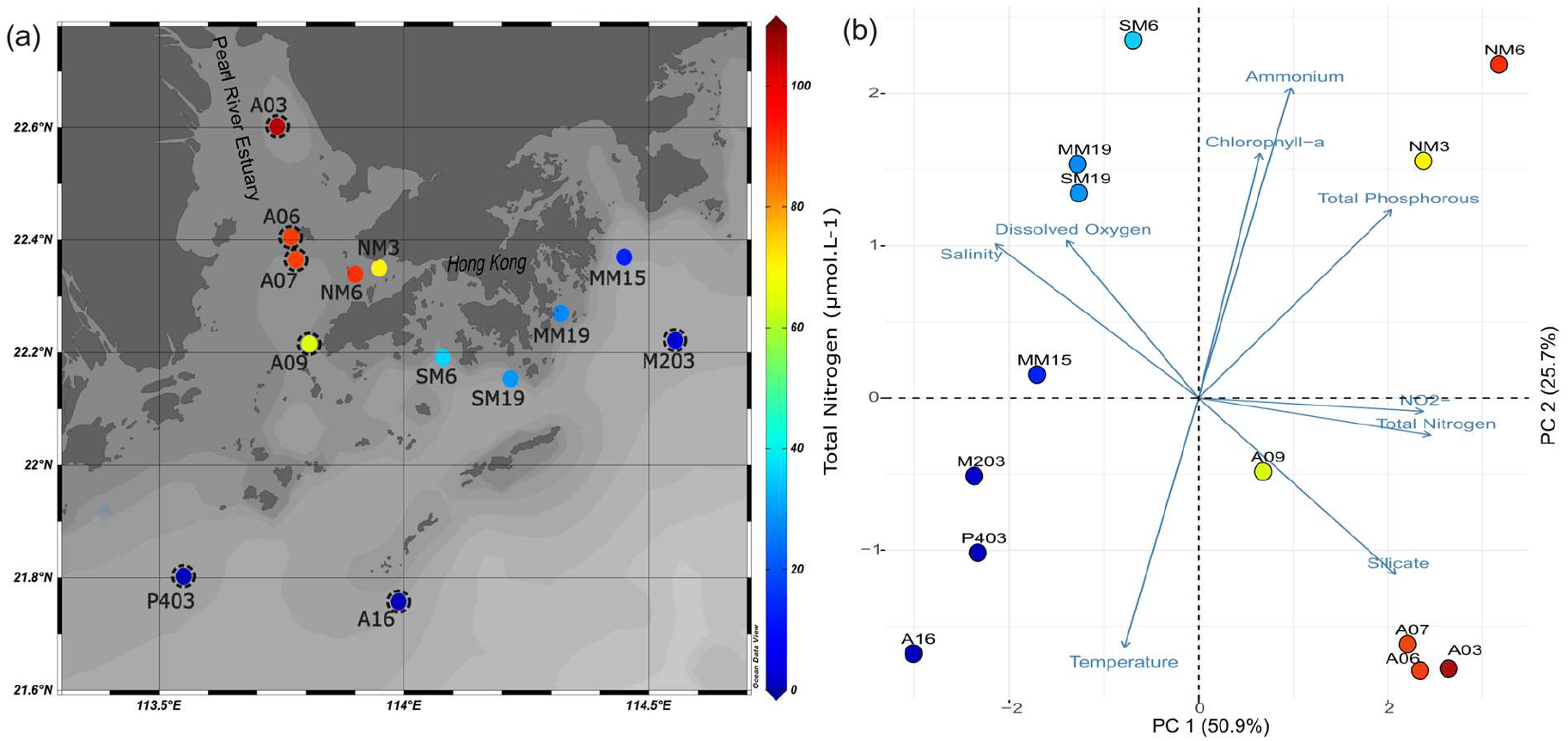
Environmental condition of the sampling sites. (a) Location of the zooplankton sampling and DOM released incubation along the Pearl River Estuary and adjacent coastal waters. (b) Principal component analysis (PCA) biplot of environmental parameters. The stations for zooplankton derived DOM release incubation (summer 2024 cruise) were indicated by dashed circles. The colours dots denote nutrient level (total nitrogen) while the arrow length represents the contributions of each environmental parameter toward the variance of principal component.

We conducted a principal component analysis (PCA) on measured environmental parameters to assess the environmental heterogeneity across the sampling stations (Fig. 1 (b)). The first two principal components (PC1 and PC2) collectively explained 76.6% of the total variance across sampling sites. PC1 was primarily driven by nutrient concentrations, including total nitrogen, nitrite (NO_2_^−^), and salinity, while PC2 reflected variations in ammonium, total phosphorus, and temperature. Given the distinct nutrient gradients observed, we classified regions using trophic state indices, revealing a clear separation into mesotrophic (blue dots) and eutrophic (yellow to red dots) regions (Fig. 1a).

DOM analysis identified a total of 4,945 molecular formulas, with an average of 900 formulas for each sample of zooplankton-derived DOM. These molecules were categorized into different classes, including lipids, proteins, amino sugars, carbohydrates, unsaturated hydrocarbons, lignin, tannins, and condensed hydrocarbon-like compounds, based on their H/C and O/C ratios threshold, as shown in a Van Krevelen diagram (Fig. 2a). Notably, nearly half (48.8%) of the DOM molecules were common to both eutrophic and mesotrophic zooplankton communities, while 30.8% and 20.5% were exclusive to eutrophic and mesotrophic systems, respectively (Fig. 2b). This overlap suggests a core set of metabolic byproducts shared across nutrient regimes, with unique molecular signatures may reflecting environment-specific physiological processes^33^.

**Fig 2.**
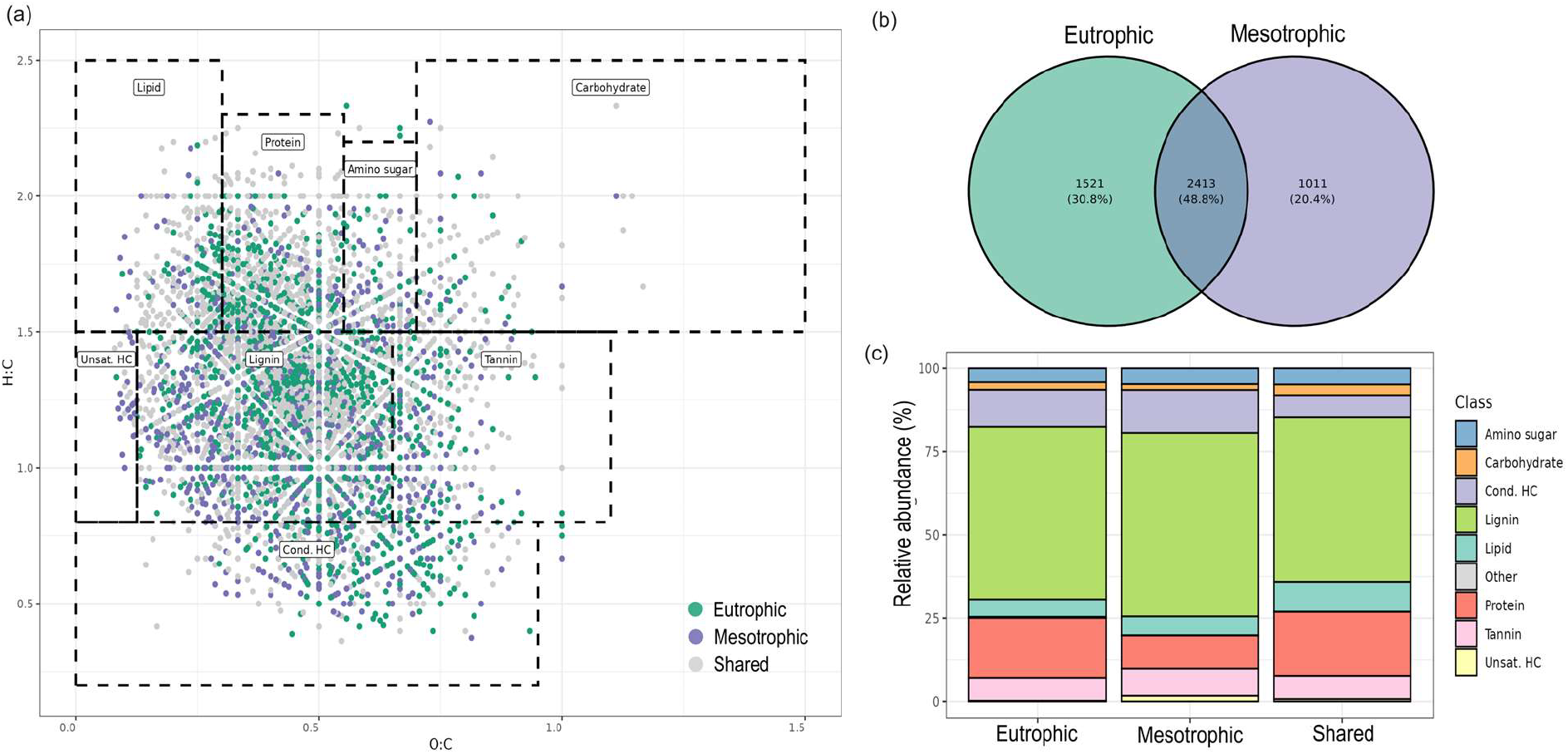
Distributions of zooplankton derived DOM molecules. (a) Van Kravelen diagram of the DOM molecules with the colours representing the molecule presences in which region trophic conditions. (b) Venn diagram of molecular formula numbers of presences in each region (mesotrophic, eutrophic, and shared). (c) Relative abundance of DOM compound like composition which uniquely presence in eutrophic, mesotrophic, and shared in both regions.

Lignin-like compounds dominated the DOM pool in all regions (55-62%), followed by protein-like (12-20%), condensed hydrocarbon-like (5-15%), and tannin-like (~5%) molecules (Fig. 2c). The comparative analysis revealed striking spatial differences in zooplankton derived DOM molecular composition. Mesotrophic zooplankton-derived DOM exhibited significantly higher proportions of lipid-like and unsaturated hydrocarbon-like compounds compared to eutrophic systems, whereas protein-like molecules were enriched in eutrophic DOM (Fig. 3a). These patterns align with the nutrient gradients where protein-like DOM in nutrient-rich systems likely reflects elevated nitrogen and phosphorous availability in the food sources (phytoplankton) and rapid zooplankton growth, driving nitrogen and phosphorous contained DOM release. Conversely, lipid- and unsaturated hydrocarbon-like compounds in mesotrophic regions may arise from lipid catabolism or structural tissue turnover under resource-limited conditions, where zooplankton rely on stored energy reserves.

**Fig 3.**
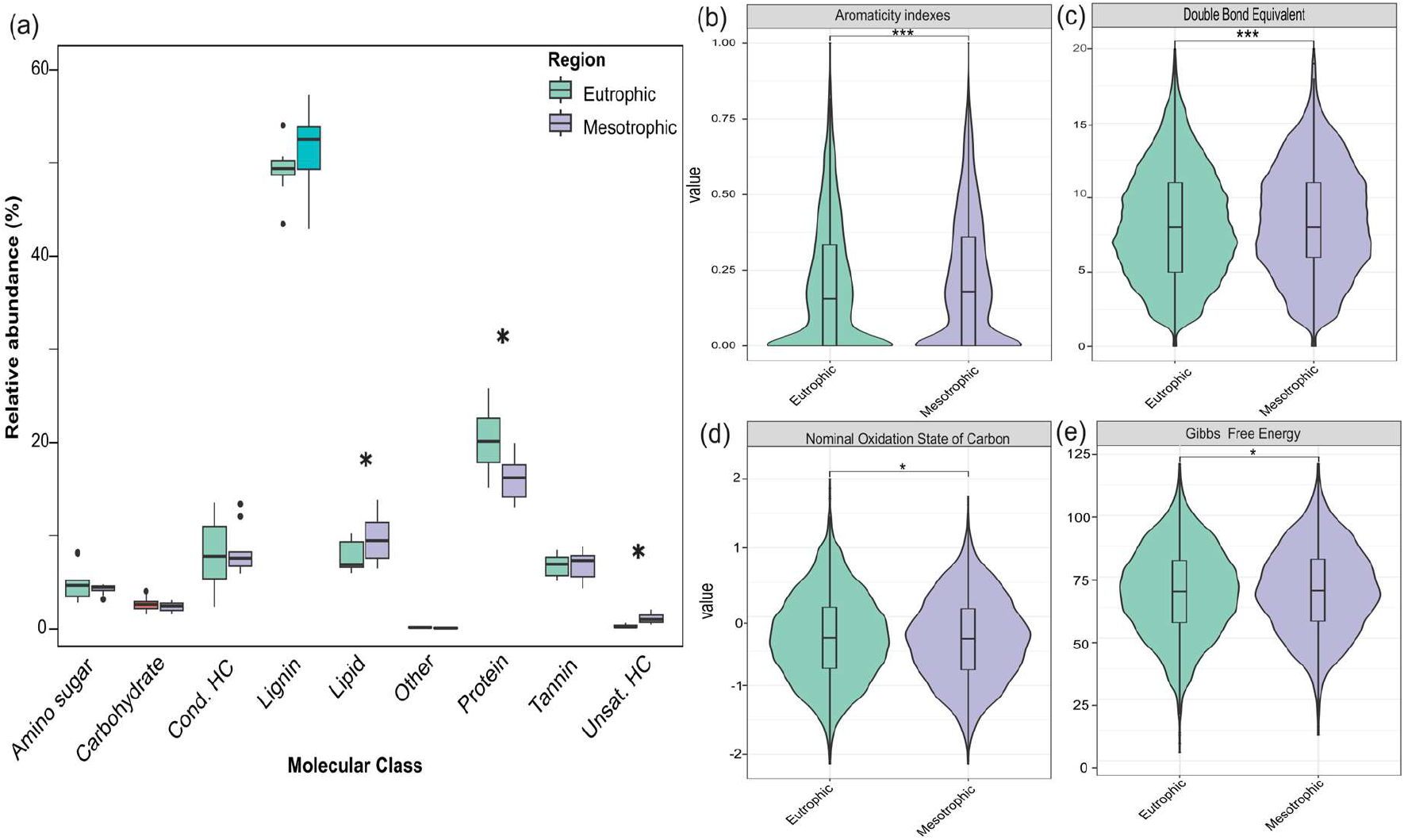
Comparison of zooplankton derived DOM molecules properties between eutrophic and mesotrophic. (a) Relative abundance (%) of DOM molecular classes with the clear difference’s molecule classes marked by asterisk. DOM chemical traits (b) Aromaticity indexes, (c) double bond equivalent, (d) Nominal Oxidation State of Carbon, and (e) Gibbs Free Energy. The colours represent the eutrophic and mesotrophic regions.

Molecular traits further underscored spatial divergence. Mesotrophic DOM displayed significantly higher modified aromaticity indices and double bond equivalents (p < 0.001; Fig. 3b–c), indicative of structurally complex, aromatic, and unsaturated molecules. Such compounds are typically recalcitrant, aligning with relatively lower Gibbs free energy values (Fig. 3e), which suggest reduced bioavailability and slower microbial degradation rates. In contrast, eutrophic DOM showed a higher nominal oxidation state of carbon (NOSC; Fig. 3d), indicating greater oxidative potential and labile substrates.

### Zooplankton community gene expression profiles highly associated with environmental factors

To investigate physiological variability in zooplankton communities across sampling regions, we conducted a meta-transcriptomic analysis. Taxonomic classification revealed 20 zooplankton classes (Fig. S1), with the top 20 classes representing ~80% of total relative abundance. Dominant taxa included *Calanoida, Harpacticoida*, and *Siphonostomatoida*, consistent across all regions. Functional annotation identified 12,663 transcripts with Kyoto Encyclopaedia of Genes and Genomes (KEGG) orthologs (KOs).

To resolve metabolic heterogeneity in zooplankton communities, we analyzed KEGG-annotated gene functional profiles and their association with environmental variables using Weighted Gene Co-Expression Network Analysis (WGCNA; Fig. 4a). Principal Component Analysis (PCA) of environmental parameters revealed nutrient concentrations (e.g., total phosphorus, ammonium) as primary drivers of variance, prompting focused analysis of nutrient-associated gene modules. WGCNA identified two gene modules highly associated with nutrient concentrations, module 4 (2,096 KOs) exhibited positive correlations with phosphorus and ammonium concentrations and module 7 (341 KOs) showed inverse relationships (Fig. 4a).

**Fig 4.**
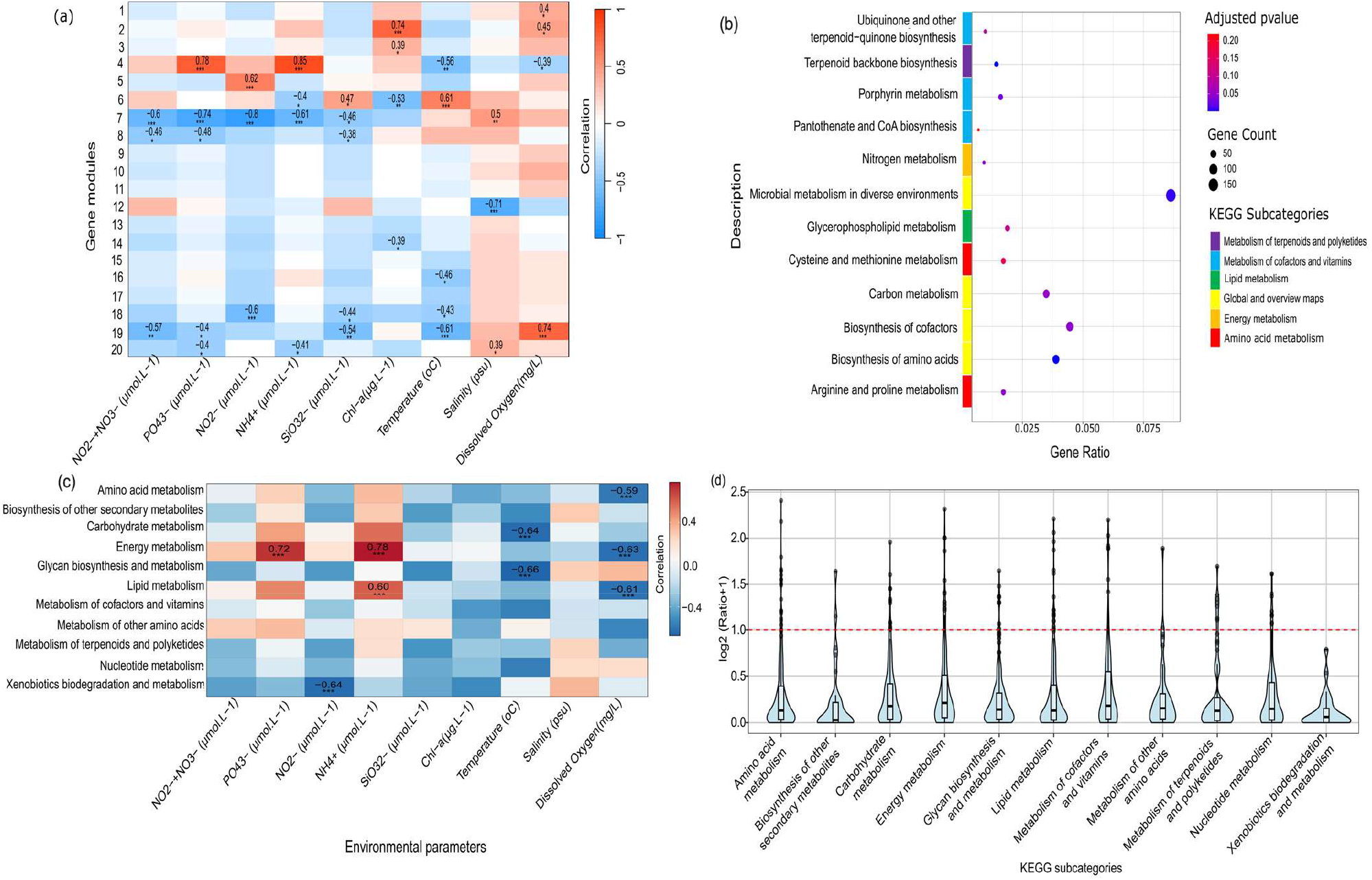
(a) Weighted gene co-expression network analysis illustrates the correlation between environmental parameters (x-axis) and groups of similarly expressed genes, assigned to gene modules (ME) (y-axis). The top numbers in each cell represent Pearson’s correlation coefficients, while the bottom numbers indicate the p-values from the correlation test. The color of each cell reflects the strength of the correlation between the ME and environmental parameters, with red indicating a positive correlation and blue indicating a negative correlation. (b) Dot plot of KEGG enrichment analysis for module 4. Enrichment is calculated based on the number of KOs overrepresented in KEGG pathways compared to all annotated KOs. Significance is determined using false discovery rate-adjusted p-values of <0.05, with darker colors indicating higher significance. The gene count represents the number of KOs detected in each pathway, and different color annotations correspond to KEGG subcategories. (c) Correlation between environmental parameters (x-axis) and KEGG metabolism subcategories (y-axis). The top numbers in each cell represent Pearson’s correlation coefficients, while the bottom asterisk indicates the significant p value (* = p value < 0.05, ** = p value < 0.01, and *** = p value < 0.001) from the correlation test. (d) Ratios of KEGG subcategory profiles between eutrophic and mesotrophic nutrient regions. The red dashed line indicates a ratio of 1; values below 1 signify lower expression of metabolic gene functions in eutrophic regions relative to mesotrophic regions, while values above 1 indicate the opposite.

Gene enrichment analysis of these modules revealed distinct transcriptional strategies tied to metabolic adaptation. The module 4 associated positively with nutrient level, exhibited enrichment in pathways associated with amino acid metabolism, energy production, cofactor and vitamin biosynthesis, and terpenoid and polyketide metabolism (Fig. 4b). This profile suggests a metabolic emphasis on biosynthesis and energy-intensive processes, consistent with environments nutrients level which support growth and compound synthesis. In contrast, the module 7 inversely linked to nutrient levels, displayed some enrichment for pathways such as retinol metabolism, drug metabolism–cytochrome P450 (both subsets of cofactor/vitamin metabolism), and linoleic acid metabolism (lipid metabolism). The latter may reflect zooplankton lipid metabolism or detoxification processes under different trophic conditions, where metabolic activity shifts toward conservation rather than biosynthesis. These findings highlight how zooplankton communities dynamically modulate core metabolic pathways— such as energy optimization under nutrient scarcity—in response to shifting trophic conditions. The results further underscore nutrient availability as a key driver of transcriptional plasticity, shaping metabolic resource allocation strategies across environmental gradients which potentially influence the physiology of zooplankton and their derived DOM molecules characteristics.

To further understand the potential zooplankton metabolic processes variabilities among different environmental conditions, we look at all the genes annotated to KEGG subcategories metabolism function. Notably, energy metabolism genes correlated positively with total phosphorus (PO_4_^3−^) and ammonium (NH_4_^+^) (Fig. 4c), indicating that zooplankton in eutrophic (nutrient-rich) environments may prioritize energy (ATP) generation, which potentially elevating DOM excretion rates of phosphorus and nitrogen-rich compounds (e.g., phospholipids, nucleic acid derivatives). Similarly, the positive association between lipid metabolism genes and NH_4_^+^ suggests nitrogen availability enhances lipid turnover in zooplankton, which will potentially increase the release of lipid-derived DOM (e.g., fatty acids, sterols). Conversely, negative correlations between dissolved oxygen (DO) and amino acid as well as energy metabolism genes imply oxygen level affect respiration and protein catabolism in zooplankton (Fig. 4c). This may reduce the release of proteinaceous DOM (e.g., amino acids, peptides) while favouring alternative pathways (e.g., anaerobic fermentation), altering DOM composition to include relatively more refractory metabolites.

To compare metabolic processes across trophic gradients, we also quantified the relative expression ratios of KEGG-annotated metabolism genes between zooplankton communities in eutrophic and mesotrophic environments. Strikingly, zooplankton in eutrophic regions exhibited reduced expression of metabolism-related genes compared to their mesotrophic counterparts (Fig. 4d). This pattern can reflect the observed physiological heterogeneity in zooplankton communities, where mesotrophic populations—facing relatively nutrient scarcity—appear to upregulate metabolic processes critical for nutrient acquisition, stoichiometric homeostasis, and survival under resource constraints. Specifically, heightened metabolic activity in mesotrophic systems likely reflects acclimatization strategies to compensate for limited nutrient availability^24,34^, such as enhanced biosynthesis of cofactors, energy-efficient substrate utilization, and recycling of essential biomolecules. Conversely, the lower metabolic gene expression in eutrophic zooplankton suggests reduced pressure to optimize nutrient uptake or balance elemental ratios, as abundant resources may alleviate the need for stringent metabolic regulation. These findings underscore how nutrient level heterogeneity may governs transcriptional pattern in metabolic pathways, with nutrient-poor environments driving physiological processes that prioritize resource optimization overgrowth.

### Zooplankton DOM molecules chemical properties highly correlated with metabolic gene functions

To investigate the molecular mechanism behind the compositional and functional properties variability of zooplankton-derived (DOM), we conducted Spearman correlation analyses between DOM molecular properties and metabolic gene functions. Our results demonstrate significant linkages between specific DOM chemical traits and zooplankton metabolic pathways (Fig. 5a). Specifically, unsaturated hydrocarbons showed positive correlations with amino acid and terpenoid/polyketides metabolism-related gene functions, while protein-like DOM molecules exhibited negative associations with carbohydrate, energy, and lipid metabolism genes. Meanwhile Lipid-like molecules demonstrated negative correlations with both energy metabolism and glycan biosynthesis/metabolism pathways. Furthermore, condensed hydrocarbon-like molecules were inversely related to terpenoid and polyketide metabolism genes. In contrast, carbohydrate and amino sugar-like compounds displayed positive correlations with energy metabolism and lipid metabolism genes, respectively.

**Fig 5.**
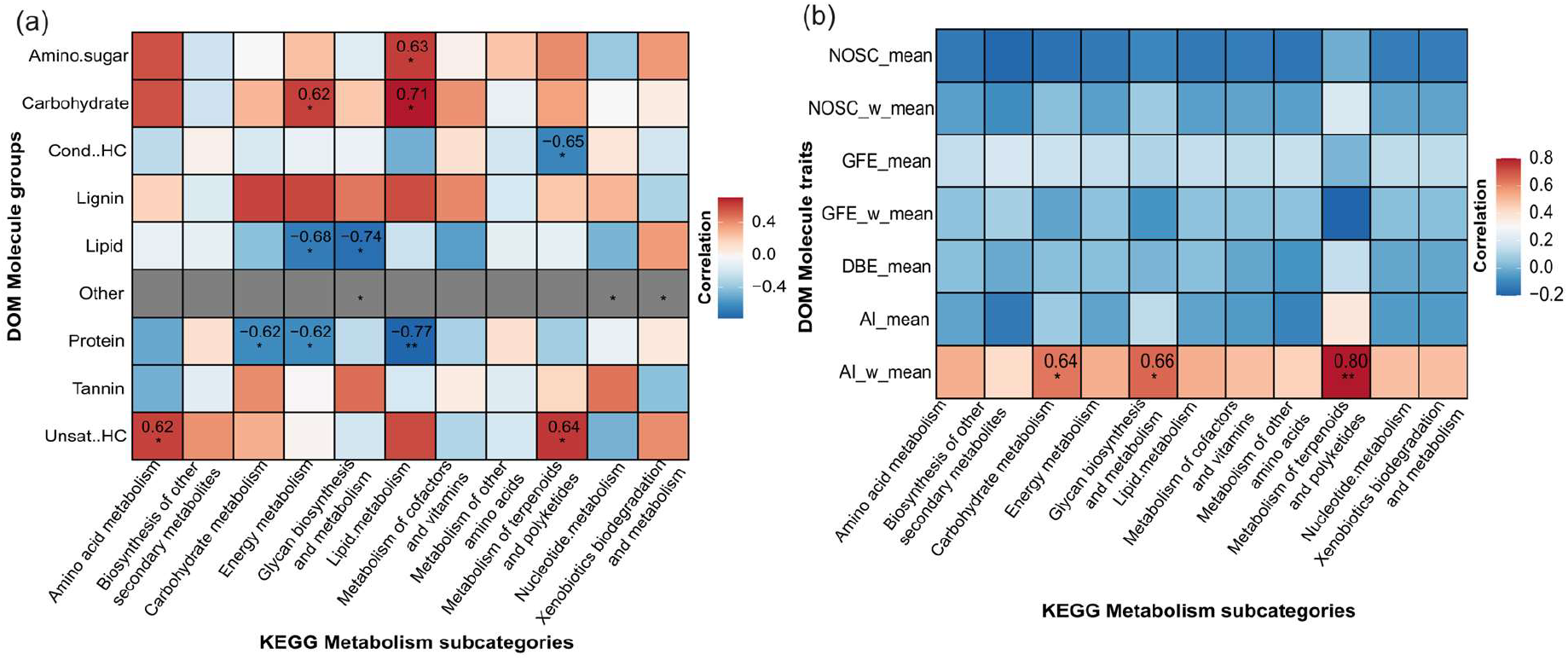
Association between DOM chemical properties with zooplankton gene functions. Spearman’s rank correlations between zooplankton’s KEGG metabolism subcategories and (a) DOM molecule groups, (b) DOM molecule traits. Significant correlations (* = p value < 0.05, ** = p value < 0.01) are indicated by the asterisks with the value inside the boxes represent correlation coefficient.

Notably, we also observed significant positive correlations between the weighted average modified aromaticity index (AI_w_mean) and three metabolic pathways: carbohydrate metabolism, glycan biosynthesis and metabolism, and terpenoid/polyketide metabolism (Fig. 5b). The observed patterns indicate that specific metabolic processes may selectively influence the chemical properties of excreted DOM, particularly its aromatic content and molecular diversity. These findings suggest that zooplankton metabolic activity plays a crucial role in determining the molecular composition and aromaticity characteristics of DOM they released into the surrounding environment.

### Implications for Nutrient Cycling and DOM Dynamics

While prior studies have extensively documented organismal physiological responses to environmental variabilities, the ecosystem-level consequences of such plasticity remain poorly resolved. Here, we bridge this gap by linking zooplankton-derived dissolved organic matter (DOM) molecular traits to transcriptional activity, revealing how metabolic gene functions shape DOM chemical properties and its role in coastal biogeochemical cycles.

Our findings demonstrate that spatial variability in zooplankton-derived DOM composition strongly correlated with transcriptional heterogeneity across sampling stations. Strong correlations between DOM molecular classes—particularly protein-like (labile, nitrogen-rich), lipid-like (semi-labile, carbon-dense), and unsaturated hydrocarbon-like (potentially more recalcitrant, reduced-carbon) compounds—and zooplankton metabolic gene expression profiles suggest that physiological responses to environmental gradients directly modulate DOM stoichiometry and bioavailability (Fig. 5a). This regulatory mechanism holds critical implications for coastal ecosystems as DOM molecular composition governs the bioavailability of bound macronutrients (e.g., nitrogen, phosphorus), which regulate microbial and phytoplankton productivity^35^. Higher or lower content of N and P within the zooplankton derived DOM compound will subsequently affect the availability of N and P in their living habitat^36^. For instance, protein-like DOM, characterized by high nitrogen content and low C:N ratios, exhibited the highest inferred bioavailability. Its production correlated negatively with carbohydrate, energy, and lipid metabolism genes, suggesting preferential resource allocation to nitrogen-rich DOM under nutrient stress. These compounds likely fuel rapid microbial assimilation and short-term nutrient remineralization, enhancing localized nitrogen/phosphorus availability. Whereas lipid-like DOM showed intermediate bioavailability, reflecting its hydrophobic properties and higher C:N ratios. Its abundance correlated negatively with energy metabolism and glycan biosynthesis pathways, implying reduced allocation in lipid content under high-nutrient conditions. Unsaturated hydrocarbon-like DOM, linked positively to amino acid and terpenoid/polyketide metabolism genes, displayed the lowest bioavailability due to structural complexity (e.g., conjugated double bonds) and enzymatic recalcitrance. These compounds may accumulate as semi-refractory carbon pools, with degradation energetics governed by their elemental composition. For instance, DOM with higher oxygen-to-carbon (O:C) ratios (e.g., protein-like molecules) requires less energy to degrade, promoting microbial growth, while low O:C compounds (e.g., unsaturated hydrocarbons) increase respiration costs and limit carbon use efficiency^37^.

Notably, zooplankton metabolic processes—particularly amino acid, carbohydrate, and terpenoid/polyketide metabolism—emerged as potential key drivers of DOM aromaticity (Fig. 5 b). This was shown by the high correlation between zooplankton metabolism gene functions activity level and the modified aromaticity indexes of the derived DOM, suggesting that zooplankton metabolism processes potentially facilitate the transformation of the released DOM into more or less aromatic compounds depending on the condition of their living environment. The strong correlation between transcriptional activity of these pathways and DOM modified aromaticity indices (AImod) implies a mechanistic link such as upregulated terpenoid/polyketide metabolism under mesotrophic conditions (relatively low nutrients) may enhance biosynthesis of aromatic secondary metabolites, increasing DOM aromaticity. Conversely, suppressed lipid or carbohydrate metabolism in eutrophic systems could favor less recalcitrant DOM production. This transcriptional tuning of DOM aromaticity has cascading effects as aromatic compounds exhibit greater biogeochemical recalcitrance due to stabilized conjugated π-electron systems^38^, potentially extending DOM residence times in the water column and altering carbon export fluxes.

## Conclusion

We integrated mass spectra and meta transcriptomics data to characterize the association between zooplankton derived DOM molecular properties and zooplankton physiological processes, and applied gene-centric analysis to elucidate the zooplankton metabolic potential that potentially shapes DOM heterogeneity. DOM chemical properties were significantly and broadly correlated with metabolism profiles of the zooplankton. These results provide evidence of how zooplankton active roles—not only intermediaries—in DOM cycling by modulating derived DOM molecules composition and aromaticity signatures through their metabolic plasticity which is strongly driven by the environmental parameters such as nutrient level.

## Methods

### Zooplankton field sampling

We collected zooplankton samples during summer 2023 (August) and 2024 (July) cruises in Pearl River Estuary-Hong Kong coastal waters from the surface (2 m depth, neustonic layer) layers using a custom-made plankton pump equipped with a 150 μm mesh sieve. The zooplankton samples were collected by deploying a plankton pump for 15 minutes with pumping rate 105 m^3^/h. The sampling sites were selected based on the long-term monitoring stations of the Environmental Protection Department Hong Kong Monitoring Program for the Summer 2023 and Earth HK project Cruise stations for Summer 2024.We categorized the sampling sites into two distinct physicochemical zones (Fig. 1a) — mesotrophic and eutrophic — based on the TRIX trophic index, a well-established metric for evaluating aquatic trophic status^39–41^.

### Zooplankton derived dissolved organic matters release experiment

We conducted incubation experiments to evaluate the release of dissolved organic matter (DOM) by zooplankton during the Summer Cruise 2024. Initially, the collected zooplankton samples were rinsed with 0.22 μm-filtered ambient seawater. Subsequently, we incubated the zooplankton in a 2-liter Watson bottle containing 1 liter of 0.22 μm-filtered ambient seawater. The control treatment consisted of 0.22 μm-filtered ambient seawater without zooplankton. The incubation lasted for eight hours in the dark at in-situ temperature. Following the incubation period, we separated the zooplankton from the released DOM using a 0.22 μm Millipore membrane filter. The collected DOM was preserved at −20°C. This incubation experiment was conducted in triplicate to ensure the reliability of the results.

### Zooplankton RNA preservation

We performed filtration using 10 µm nylon membrane to separate the collected zooplankton from both 2023 and 2024 summer cruises samples from the seawater. The zooplankton-containing filters were then preserved in Invitrogen™ RNAlater™ Stabilization Solution and stored at −80°C until processing for RNA extraction in the laboratory. Concurrently, water quality parameters including temperature (°C), salinity (PSU), and dissolved oxygen (%) were measured at each sampling station using a multi-parameter CTD-DO prior to zooplankton sampling. Additionally, nutrient concentration such as total nitrogen (mmol.m^-3^), total phosphorus (mmol.m^-3^), and silicate (mmol.m^-3^), and chlorophyll-a (μg.L^-1^) were also measured by laboratory analysis for the 2024 samples while for the 2023 we utilized complementary environmental data collected in the same sampling stations at the same month by the Environmental Protection Department (EPD) which are publicly available through the EPD data repository at Environmental Protection Interactive Centre : Marine Water Quality Monitoring Data.

### DOM extraction and characterization by FT-ICR-MS

We use solid-phase extraction (SPE) to extract dissolved organic matter (DOM) from the 0.22 μm-filtered incubation water samples following well established protocols^42^ with a recovery efficiency of around 40% for marine water DOC, and approximately 50% for freshwater water DOC. In summary, 500 mL of samples were filtered through GTTP filters (Millipore, 0.22 μm pore size, which were prerinsed three times with MilliQ water). The samples were then acidified to a pH of 2 using analytical-grade HCl and passed through a 1-g SPE cartridge (which had been prerinsed twice with methanol, MilliQ water, and MilliQ water at pH 2) (200 mg, Bond Elut, PPL, Agilent). The cartridges were desalted using 0.01 M HCl and dried with pure nitrogen gas. Finally, the SPE-DOM samples were eluted from the cartridge with 2 mL of 99.9% HPLC-grade methanol.

To characterize the molecular formulae of the SPE-DOM, methanol PPL extracts were analyzed with a 9.4 T Apex-ultra X FT-ICR MS (China University of Petroleum, CUP) under the negative mode with an electrospray ionization source (Bruker Apollo II). 128 scans were used in the mass range m/z 200 to 800 Da with the calibration of a series of established formula mass peaks (Data Analysis 3.4 software package)^43^. Peaks were selected when the signal/noise ratio (S/N) was greater than 4. Formula were assigned according to the formula rules^44^, allowing for elemental compositions C_1-60_H1-120O_1-40_N_0-3_S_0-3_P_0-1_ with an error range of ± 1 ppm. The annotated DOM molecules were assigned to biochemical compound categories based on the stoichiometry of C, H, and O for the following H:C and O:C ranges; lipids (0 < O:C ≤ 0.3, 1.5 ≤ H:C ≤ 2.5), unsaturated hydrocarbons (0 ≤ O:C ≤ 0.125, 0.8 ≤ H:C < 2.5), proteins (0.3 < O:C ≤ 0.55, 1.5 ≤ H:C ≤ 2.3), amino sugars (0.55 < O:C ≤ 0.7, 1.5 ≤ H:C ≤ 2.2), carbohydrates (0.7 < O:C ≤ 1.5, 1.5 ≤ H:C ≤ 2.5), lignin (0.125 < O:C ≤ 0.65, 0.8 ≤ H:C < 1.5), tannins (0.65 < O:C ≤ 1.1, 0.8 ≤ H:C < 1.5), and condensed hydrocarbons (0 ≤ O:C ≤ 0.95, 0.2 ≤ H:C < 0.8). Unnamed compounds (others) are calculated as the proportion of identified C that does not fit within the above defined H:C and O:C ranges^45^. Relative peak intensities were calculated based on the sum-normalized intensities of all assigned peaks in each sample^46^.

### RNA extraction, library preparation, and sequencing

The collected zooplankton samples preserved in RNAlater were used for zooplankton total RNA extraction by using FastPure® Cell/Tissue Total RNA Isolation Kit V2 following the manufacturer’s protocol. For the purpose of RNA quantity and quality checking, Qubit, Biodrop, and Gel Electrophoresis were used prior to library creation and sequencing. In the RNA sequencing step, each sample was processed by isolating mRNA from one microgram of total RNA through poly-T oligo-attached magnetic bead selection. Subsequently, the mRNAs were reverse transcribed into cDNAs to construct RNA sequencing libraries with average insert sizes ranging from 250 to 300 bp, utilizing TruSeq™ RNA Sample Prep Kits (Illumina) in accordance with the manufacturer’s guidelines. To ensure robustness and account for biological variability, two to three biological replicates were included for each sample, resulting in a total of 34 libraries. The metatranscriptomes were subjected to pair-end sequencing, producing 150 bp reads using the HiSeq™ 2000 sequencing platform (Illumina) following standard protocols.

### Meta transcriptome de novo assembly, transcript abundance estimation, and functional annotation

The raw RNA sequence data quality was evaluated using FastQC Version 0.11.5^47^, followed by quality trimming of the raw reads for poor quality bases and RNA-Seq adapters using Trimmomatic. The resulting paired-end clean sequence from all 34 libraries were simultaneously utilized to construct the meta-assembly using Trinity software v2.15.2 with default setting^48^. Raw reads were quality-filtered and assembled de novo, yielding 12,975,434 contigs. The assembled contigs then being used for open reading frame prediction by using transdecoder against pfam database. Subsequently, the reading alignment and transcript abundance estimation was performed by using Salmon-based approach. In brief, the reads are first aligned to the assembled contigs using Bowtie2. The output BAM file is then used as the input for Salmon tool. Normalization of transcript per million (TPM) was performed based on library sizes using the “calcNormFactors (method = “TMM”)” function to ensure accurate quantification of gene expression levels across the samples^49^.

To obtain functional annotations for the assembled contigs, we utilized the Trinotate pipeline^50^. This process also needs protein sequence as one of the inputs for the functional annotations, consequently we used Transdecoder software packaged with Trinity to predict the protein translations from the contigs. Subsequently, the unigenes obtained were queried against several public databases, including SwissProt, Pfam-A, and Eggnog^51^, with an E-value threshold of less than 1E^-5^ using the Trinotate pipeline. In this annotation process, BLAST 2.9.0+ was employed for gene assignments, using diamond BLASTp for protein queries and diamond BLASTx for nucleotide queries^52^. Both query sets were compared against the Uniprot/SwissProt database of reference proteins^53^. To identify conserved domains within the protein models, we utilized HMMER v.3.3 and the Pfam-A database^54,55^. The results were then loaded into Trinotate’s SQL database. Trinotate used the top BLAST hits to populate the database with gene ontology (GO) and Kyoto Encyclopedia of Genes and Genomes (KEGG) terms.

### Statistical analyses and data visualization

We employed the MetaboDirect pipeline^56^ to conduct data exploration and statistical analysis, including the generation of van Krevelen diagrams, Venn diagrams, and visualizations of dissolved organic matter (DOM) relative abundance. The input data consisted of preprocessed DOM molecular formulas along with their corresponding monoisotopic peak intensities and *m/z* values. To evaluate molecular properties and potential decomposability, we computed several thermodynamic and molecular indices based on each peak’s elemental composition (equations provided in Supplementary Table 2). These indices included the Nominal Oxidation State of Carbon (NOSC), which describes the average carbon oxidation state of the DOM molecules; Gibbs free energy (ΔG°C-ox or GFE), indicating the thermodynamic favorability of degradation^57^;the Modified Aromaticity Index (AImod), reflecting aromaticity and carbon double-bond density^58^;and the double bond equivalent (DBE), quantifying molecular unsaturation as a proxy for DOM aromatic structures^59^.

During data exploration, MetaboDirect pipeline generated van Krevelen diagrams^60^ to visualize molecular formula distributions, along with violin plots of the calculated thermodynamic indices. Statistical differences between eutrophic and mesotrophic DOM release were assessed using Tukey post hoc tests, while bar plots summarized molecular and elemental compositions across different regions. Additionally, pairwise comparisons were performed to identify unique and shared molecules between mesotrophic and eutrophic groups, employing van Krevelen diagrams, Venn diagrams, and stacked bar plots for visualization.

To investigate zooplankton gene functional profiles and their relationship with environmental variables, we conducted a weighted gene co-expression network analysis (WGCNA) using the WGCNA package (version 1.72-1) in R^60^. The analysis utilized a trimmed mean of M-values (TMM) normalized expression matrix of Kyoto Encyclopedia of Genes and Genomes (KEGG)-annotated transcripts as the input to identify gene modules exhibiting strong co-expression patterns. Network construction employed an unsigned topological overlap matrix (TOM), with module detection based on TOM-derived dissimilarity measures (1-TOM). Parameters were optimized for biological network properties, including a soft threshold power (β) of 30 to satisfy scale-free topology criteria, minimum module size of 70 genes, and a merge cut height of 0.4^22^. Module eigengenes were computed and correlated with both environmental parameters and zooplankton taxonomic group relative abundances. Gene enrichment analysis was performed using the clusterProfiler package in R to identify metabolic pathways associated with significant module eigengenes. Resulting p-values were adjusted for multiple comparisons using the Benjamini-Hochberg false discovery rate (FDR) correction^22,61^.

We employed Spearman rank correlation analysis to examine relationships between KEGG metabolic subcategories and both environmental factors and DOM chemical characteristics, including molecular class compositions and molecular traits. To compare zooplankton metabolic processes between trophic states, we calculated eutrophic-to-mesotrophic ratios of total transcripts per million (TPM) values for each KEGG metabolic subcategory.

## Acknowledgements

The work described in this paper was substantially supported by the funding from Center for Ocean Research in Hong Kong and Macau (CORE), the joint research centre for ocean research between Qingdao National Laboratory for Marine Science and Technology (QNLM) and Hong Kong University of Science and Technology (HKUST). This study is also partially supported by Research Grants Council of the Hong Kong Special Administrative Region, China (project reference no. AoE/P-601/23-N). We thank J. Poon, L. Yuanmei, Y.yan for assistance with the fieldwork. We also thank M. Ye for lab works and J. Li for the bioinformatics assistance.

